# GraTools, a user-friendly tool for exploring and manipulating pangenome variation graphs

**DOI:** 10.64898/2025.12.01.691558

**Authors:** Sebastien Ravel, Nina Marthe, Camille Carrette, Mourdas Mohamed, Francois Sabot, Christine Tranchant-Dubreuil

## Abstract

**Background:** Pangenome variation graphs (PVGs), which represent genomic diversity through multiple genomes alignment, are powerful tools for studying genomic variations in populations. However, current tools often lack integration, efficiency, or require format conversions, to use them, hindering their usability.

**Results:** Here, we introduce GraTools, a member of the GraSuite [1], and a fast and user-friendly command-line tool for manipulating PVGs using the original GFA file. After a one-time graph import, GraTools enables rapid subgraph extraction, FASTA sequence retrieval, and comprehensive analyses, including core/dispensable genome ratio calculation or group-specific segment identification. The import step results in conversion in standard data formats (BAM/BED), enabling the reuse of well-optimized existing tools, allowing an efficient storage and the querying of the PVGs large complex data structures. Scalability is ensured by a modular architecture supporting parallel processing and asynchronous I/O operations. GraTools supports coordinates defined on both the primary reference as well as from alternative genomes within the graph without re-import, and its outputs can be easily visualized or manipulated using external tools. Using an Asian rice pangenome graph (13 accessions), we demonstrate its ability to easily extract subgraphs, compute depth statistics, and identify subspecies-specific segments. An intuitive command-line interface, a real-time execution feedback and a detailed logging system make this tool suitable for a wide range of applications, from population genetics to breeding and genomic medicine, for both biologists and bioinformaticians.

**Conclusions:** Through its unified graph manipulation interface, GraTools offers an interesting alternative to the few existing tools for manipulating PVGs, facilitating rapid, efficient and flexible downstream analyses. It is available as an open-source tool (GNU GPLv3), with its documentation available at https://gratools.readthedocs.io.

## 1 Introduction

Nowadays, more and more studies on diversity, agronomy, health or evolution, at the species or genera level, focus on the concept of pangenome. This approach, that considers the whole genomic content of a group of individuals [2, 3] and not of a single individual, allows to better identify variations and thus the evolution and functional role of sequences (e.g. Sub1A gene in rice [4]). While many types of pangenome approaches exist [2, 5], in higher eukaryotes and for species-level analyses, most of the recent studies focus on pangenome variation graphs (PVGs) for representing the pangenomic sequence variations. These PVGs, generally built from a MSA-like approach [6, 7], are directed graphs, in which nodes (or segments or fragments) represent sequences of varying lengths from at least one individual, and edges (or links) indicate adjacency between two sequences along at least one haplotype (or path or walk) in the PVG[8, 9].

The most currently used format for storing PVGs is the GFAv1.1 (Graphical Fragment Assembly [10]), a text tabular-based format structured in S, L and W/P lines, for segments, links and walks/paths, respectively. Using the W/P lines, one can reconstruct the genomic sequence of an embedded individual through the sequences of the nodes within the S lines. Other formats exist for storing PVGs, such as the rGFA or GFAv2.0, but are lacking some information (no P/W lines in rGFA e.g.), or are not widely spread in the community (for GFA2.0).

Many PVGs construction tools provide a graph in GFAv1.1 format, such as MiniGraph/Cactus [7], PGGB [6] or vg [8]. In addition, different tools offer functionalities for PVGs analysis, including basic statistics (gfastats [11], pancat [12]), basic manipulations (vg [8], odgi [13], gfatools [14]), subgraph extraction (pancat [12]), and retrieval of individual FASTA sequences (vg [8], odgi [13]). However, none of those tools provide an integrated suite for manipulating PVGs through a single, simple, and user-friendly command-line interface. In addition, some tools exhibit limitations in terms of execution time or overall performance for downstream analyses, or require GFA files to be converted into their own format and this new format referred for all command, or to modify the initial file outside of the tool (e.g., node renaming) prior to analysis.

Here, we propose GraTools, a member of the GraSuite [1], an intuitive and fast command-line tool for manipulating PVGs using the initial GFA file as query name (see below), and conserving node names as input and output, in a transparent way for the user. It converts the graph in an optimized data structure once and allows for instance users to extract subgraphs and fasta sequences either from a given location (genome path or graph nodes) or based on a specific composition. It also provides statistics on the global PVG and on the Core/Dispensable compartment, and can perform basic analyses on the graph topology.

We provide here some examples as use-cases of increasing complexity in order to illustrate GraTools efficiency, using an Asian rice PVG [15] as example. While GraTools is primarily designed to efficiently work with PVGs coming from MiniGraph/Cactus [7], it also partially works with the ones coming from PGGB [6] after conversion (path to walk with coordinates), using tools such as GraMer [16]. GraTools is developed in Python, installable using *pip* and an *Apptainer* or *Docker* container. The GraTools code and the supplementary scripts used here are available under the GNU GPLv3 license at https://forge.ird.fr/diade/gratools and https://data.ird.fr/dataverse/gratools, and the complete GraTools manual at https://gratools.readthedocs.io/.

## 2 Implementation

GraTools is implemented in Python 3.12+ and designed to be a robust and efficient tool for pangenomic variation graph analysis. It requires a single external dependency, BEDtools [17], which must be available in the system path.

### 2.1 Installation and manual

The tool can be installed *via* the Python Package Index (PyPI), from its source repository or using *Apptainer* and *Docker* containers. Documentation, including installation instructions, tutorials, and a detailed command reference, is hosted on *ReadTheDocs* (https://gratools.readthedocs.io).

### 2.2 One-Time Importing

All the commands of Gratools take a GFA file as input. To enable efficient subsequent graph querying, each GFA is imported once in BAM and BED format (see below), either explicitly using the gratools import command, or automatically when any other command is invoked (if no import has been previously performed). During this step, the graph is parsed and stored in two optimized formats: an indexed BAM file for segments information, and a collection of BED files containing segments positions across individual genomes. The import phase is the most computationally intensive step, typically requiring from less than one minute to more than 30 minutes depending on the PVG size (see section 3.1). Once completed, subsequent analyses are performed without re-parsing the graph, substantially reducing execution time. This internal file structure is fully transparent to the user, as said before, as the indicated input file for all the GraTools commands is always the initial GFA file.

### 2.3 Architecture and Core Components

GraTools is built on a modular, class-based architecture in which each component has a clearly defined role. A central main Gratools class orchestrates interactions with three other main classes, as illustrated in Figure 1.

**Fig. 1.**
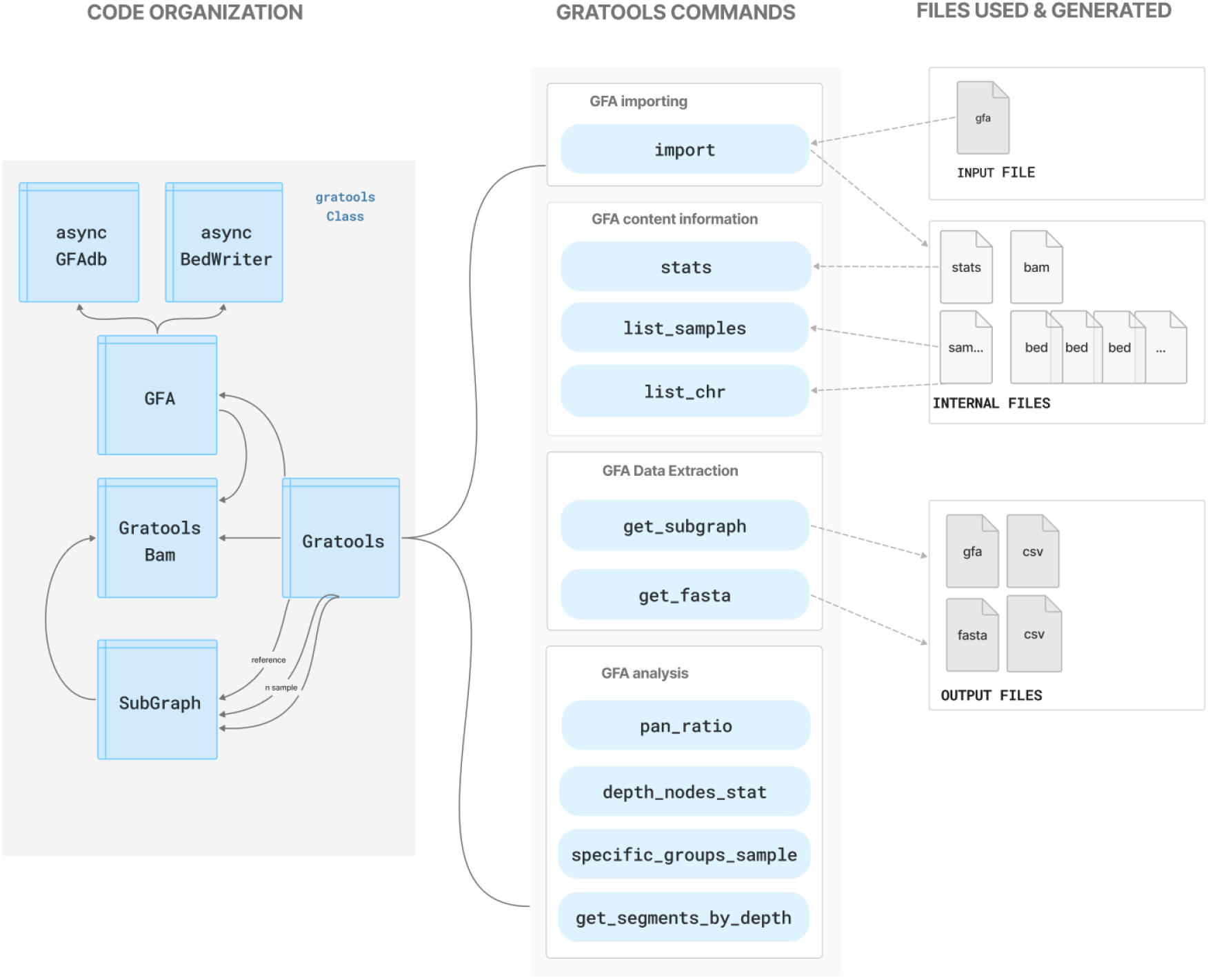
Overview of GraTools: GraTools follows a class-based architecture in which the main Gratools class coordinating interactions with the three other classes: GFA, GratoolsBam, and SubGraph. The functionalities and commands are organized into four categories: **(i) GFA importing** (import), **(ii) GFA content information** (stats, list samples, list chr), **(iii) data extraction** (get subgraph, get fasta), and **(iiii) content analysis** (pan ratio, depth nodes stat, specific groups sample, get segments by depth). The initial GFA file is used as input file, and outputs are provided in FASTA, GFA and CSV formats. Internal files (BAM and BED) are used by GraTools to speed up processing, transparently for the user.

#### Gratools Class

This class acts as the main orchestrator of all the GraTools commands, serving as the bridge between the command-line interface and the backend processing modules. This class interprets the user commands, manages input and output paths, and coordinates their execution.

#### GFA Class

This class is responsible for the initial import of the GFA file, including its parsing and processing. It reads each line and extracts the relevant information to populate the internal data structures. During this phase, it forwards path information to the AsyncBedWriter for BED files creation, and sends link information to the AsyncGfaDatabase for database indexing (optional).

#### GratoolsBam Class

The GratoolsBam class manages all interactions with the BAM file generated during the initial GFA import. It handles essential operations such as writing, indexing and reading the BAM file, as well as querying it to extract specific nodes. This class forms the basis of the computational engine for most sophisticated commands. By leveraging the internal BAM file and the pysam library [18–21] in particular, it efficiently performs complex calculations such as core/dispensable ratios, segment depth statistics, or identification of shared segments between sample groups.

#### SubGraph Class

The SubGraph class extracts user-defined genomic regions from any embedded path in a graph. It intersects query coordinates in the pre-computed BED files using pybedtools [22], identifies the corresponding nodes and paths, and quickly reconstructs the subgraph in memory. The resulting subgraph can then be exported in GFA format (with native node identifiers), or as linear FASTA format.

#### AsyncGfaDatabase and AsyncBedWriter Classes

These two classes handle the various I/O operations during the import step. AsyncBedWriter creates BED files for individual genomic paths per sample, while AsyncGfaDatabase manages an asynchronous SQLite database to store and query all link data from the GFA file (optional). Batch insertion efficiently handles millions of links during the initial parse. Once the GFA file is read, internal files are created to enable high-speed queries for graph traversal and connectivity, such as retrieving segments connected to a given node, or identifying distinct connected segments.

### 2.4 Performance and Scalability

Designed for large-scale pangenome graphs analyses, GraTools optimizes performance through asynchronous I/O, efficient data handling and parallel processing. I/O operations during GFA import are managed asynchronously through dedicated Python libraries such as asyncio, uvloop [23], aiosqlite [24], aiofiles [25], enabling simultaneous writing. After the GFA import, all PVG data are accessed through BAM and BED files, with BAM queries handled by pysam [18–21] and subsequent data manipulation on BAM or BED performed using pybedtools[22] or pandas [26, 27]. CPU-bound tasks, particularly those iterating over numerous samples, are parallelized using the Python library concurrent.futures.ProcessPoolExecutor [28], allowing the user to specify the number of cores and substantially reducing computational time for large analyses.

### 2.5 User Experience and Interactivity

A key design goal was to develop a command-line tool that is efficient, intuitive for most users, and transparent during its execution.

#### Command-Line Interface (CLI)

The CLI, built with the Click, click-extra, and cloup libraries [29–31], provides a clean, structured, and color-coded interface. Commands are grouped logically, with detailed help for each command and option, and shell completion facilitates rapid and accurate command entry.

#### Interactive Feedback

For long-running operations such as importing or parallel analysis, GraTools provides detailed, real-time visual information using the rich library [32] with multi-level progress bars that track both overall progress and individual subtasks (e.g., sample processing), giving clear visibility into execution steps progress.

#### Advanced Logging

GraTools implements a non-blocking logging system with log handling in a separate background thread, minimizing performance impact on the main application and supporting multiple verbosity levels with color-coded output.

## 3 Results

GraTools can perform a wide range of operations on PVGs from a GFA1.1 file (as described in Table 1) preferentially from MiniGraph/Cactus, many of which previously required many different tools (Table 2). However, in these other tools, most of the commands are multiples, requiring sometimes the use of another external tools, and referring to internally created files as inputs (Table 2). In addition, the node names may change between input and output. As described in Section 2.2, from the end-user perspective, in GraTools everything appears to be performed directly on the initial GFA file. The optimized internal data structures created by GraTools during the import remain hidden, and outputs in GFA format preserve the original nodes, paths and haplotype names.

**Table 1.**
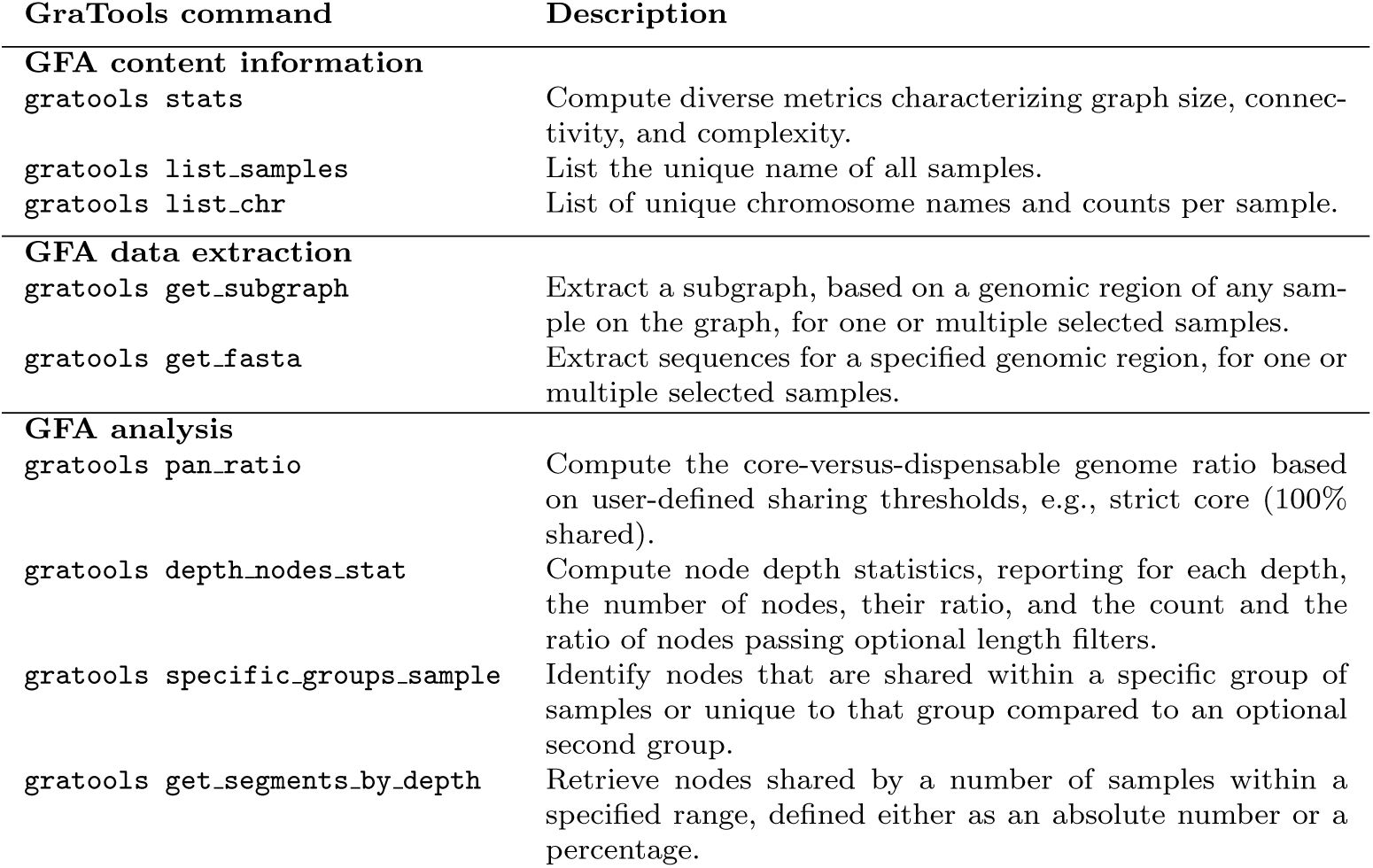
Complete list of GraTools commands.

**Table 2.**
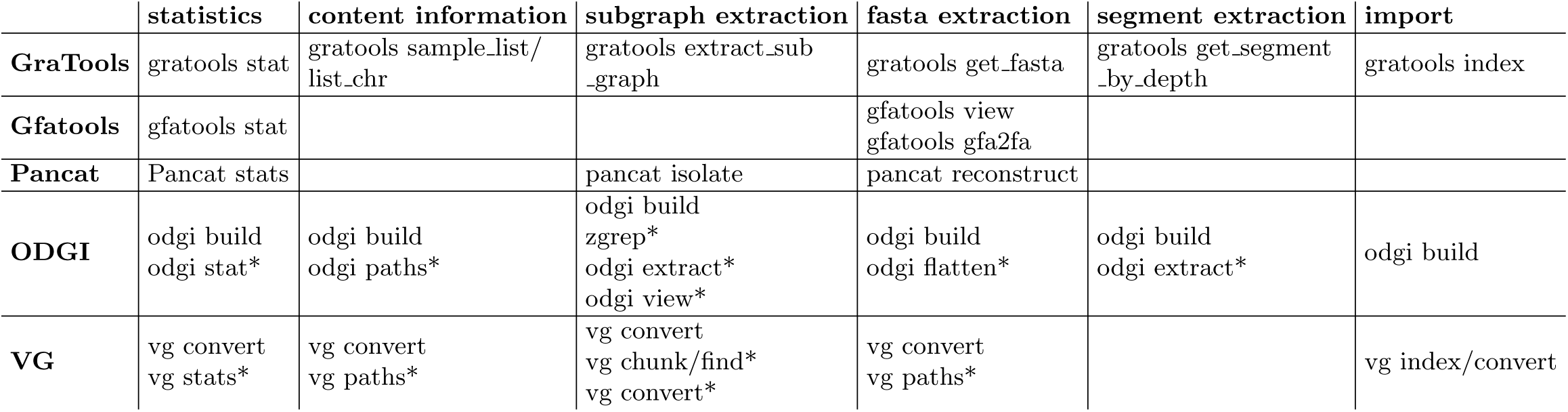
Comparative table of GraTools and other PVGs manipulation tools. Most GraTools feature are available on other tools, but the GraTools version is generally simpler and more explicit. For instance, the odgi extract command can select paths to extract, but it is node-based and not position-based, thus additional external commands (awk, grep,…) are needed to identify the desired nodes. In addition, post-import commands for odgi and vg (signaled with an asterisk) require the use the name of the newly created file or a modified one as input and not the native GFA file name.

Based on the use-cases presented below, the GraTools features can be organized into three main sections: basic description of the graph, extraction of subgraphs and fasta sequences, and analyses of graph content and genomic diversity. The usage examples presented here are mostly based on the Asian Rice graph created in [15], which includes 13 accessions from Zhou et al [33], available at https://dataverse.ird.fr/ dataverse/grannot and called **NewRiceGraph MGC.gfa.gz**. We present 3 classical analyses, each illustrating a main type analysis, and a fourth example, requiring additional commands. These examples progress from simple to more complex use cases, addressing different biological questions.

### 3.1 Benchmark

All the computations were performed on a Intel ®Xeon ®Gold 6238R CPU @ 2.20GHz, with 56 cores hyperthreading and 512 GB of RAM, running under RockyLinux 9.1 BlueOnyx. We benchmarked different types of graphs (see Table 3) under different conditions of multithreading for the import command. As shown on Table 4, the use of 10 cores improved the speed of importing without increasing the RAM usage.

**Table 3.** Information about benchmarked PVGs in GFA 1.1 format. They are all built using MiniGraph/Cactus, as in Marthe et al [15]. The number of samples and the number of chromosomes or contigs per samples are indicated.The *E. coli* and *Oryza sativa* (Asian rice) data are from Marthe et al [15], and the *Bathycoccus prasinos* ones are from Dennu et al [39].

| Graph | Pangenome size in bp | Nb of Samples | Nb of chr/sample | Reference |
| --- | --- | --- | --- | --- |
| *E. coli* | 11,630,600 | 13 | 1 | [15] |
| *B. prasinos* | 34,311,243 | 31 | 18-21 | [39] |
| *O. sativa* | 858,417,045 | 13 | 12 - 508 | [15] |

**Table 4.** Benchmark of GraTools import command on various PVGs listed in Table 3. We used different CPU number to test the multithreading efficiency. Results are in CPU time and in RAM usage (in GB, between brackets). The size of the resulting files in GB is provided.

| Graph | Pangenome size | CPU, 1 thread | CPU, 10 threads | Output files size |
| --- | --- | --- | --- | --- |
| *E. coli* | 11 Mbp | 00:00:43<br>(1.1 GB) | 00:00:30<br>(1.2 GB) | 381 MB |
| *B. prasinos* | 34 Mbp | 00:05:51<br>(8.3 GB) | 00:04:50<br>(8.3 GB) | 2.9 GB |
| *O. sativa* | 858 Mbp | 00:26:47<br>(36 GB) | 00:24:31<br>(36 GB) | 15 GB |

Compared to vg and odgi, GraTools is slightly slower when importing the Asian rice PVG (Table 5), with odgi being the fastest (8 min) and GraTools the slowest (24 min). In terms of RAM usage, GraTools import has the larger memory footprint (36 GB compared to 6 and 11 for vg and odgi, respectively). However, the GraTools import, once performed, can be used without modifications for calling regions from any of the embedded haplotypes in the graph, at the opposite of vg and odgi, where the index is strictly limited to a single ”reference” genome for extraction (see below).

**Table 5.**
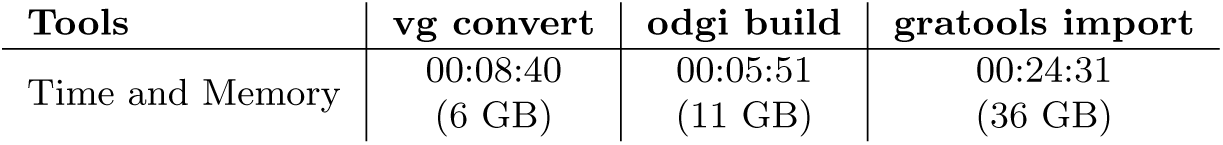
Comparison of GraTools import vs odgi build and vg convert. All tests were performed using 10 threads and on the Asian rice PVG as described in Table 3. RAM usage is provided in GB, between brackets.

| Tools | vg convert | odgi build | gratools import |
| --- | --- | --- | --- |
| Time and Memory | 00:08:40<br>(6 GB) | 00:05:51<br>(11 GB) | 00:24:31<br>(36 GB) |

### 3.2 Description of the graph content

#### 3.2.1 Obtaining basic statistics and importing

Similarly to many tools working with GFA files, GraTools proposes basic statistics about the PVG with the stats command including the number of nodes, their mean, minimal and maximal size, the mean and maximum number of output branching (using dedicated options), and the total graph size. These statistics are computed internally during the import of the initial GFA; however, they can be re-called also directly through the gratools stats command, that will be performed the import command if it has not been performed before.

Here, as a use case, we want to know which individuals are embedded in the Asian rice PVG, their chromosomes/contigs length and number, and the basic statistics on the graph. The command gratools stats --gfa NewRiceGraph MGC.gfa.gz provides the basics statistics listed in Table 6. In a simple command, we have access to the mean segment size (32.44 in the example), the average degree (2.74), the total size of the graph (858 Mb), the distribution length of nodes, etc.

**Table 6.**
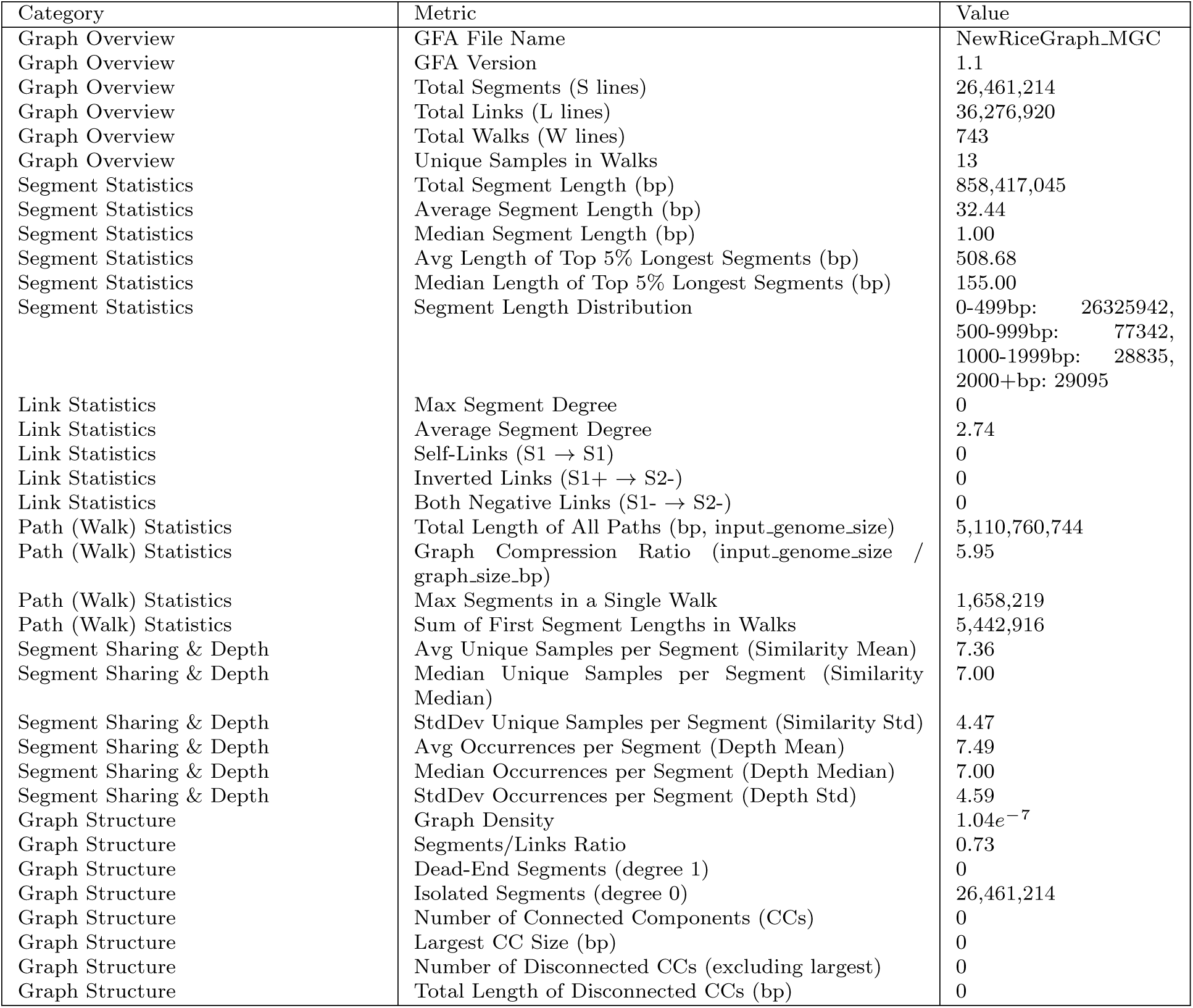
Statistics on the PVG from 13 Asian rice genomes from Zhou et al [33] and built using MGC in Marthe et al [15]. The command gratools stats can be called directly from the native graph, or after an gratools import command.

#### 3.2.2 Listing the path names and the chromosome groups

The list samples command will output the list of individual paths embedded in the graph, as shown in Table 7, while the list chr will display the chromosomes (or contigs) list organized by sample (Table 8). These features can be helpful to quickly explore a graph, to know which samples (haplotypes) were embedded in it, or what are the homologous chromosomes/contigs that the graph builder considered (if any) to be grouped. The Table 8 provides also the fragments start and stop, *i.e.* the size of the genomic sequences for each part of the chromosomes/contigs embedded in.

**Table 7.** Adapted output that list the embedded samples in the GFA from Asian rice, obtained through the gratools list samples --gfa NewRiceGraph MGC.gfa.gz command. General statistics are outputted before the table, in the terminal as in the CSV output.

| **General statistics** |
| --- |
| Total samples in GFA: 13 |
| Available Samples in GFA: NewRiceGraph_MG |
| **Sample Name** |
| AzucenaRS1 |
| IRGSP |
| Os117425RS1 |
| Os125619RS1 |
| Os125827RS1 |
| Os127518RS1 |
| Os127564RS1 |
| Os127652RS1 |
| Os127742RS1 |
| Os128077RS1 |
| Os132278RS1 |
| Os132424RS1 |
| OsIR64RS1 |

**Table 8.**
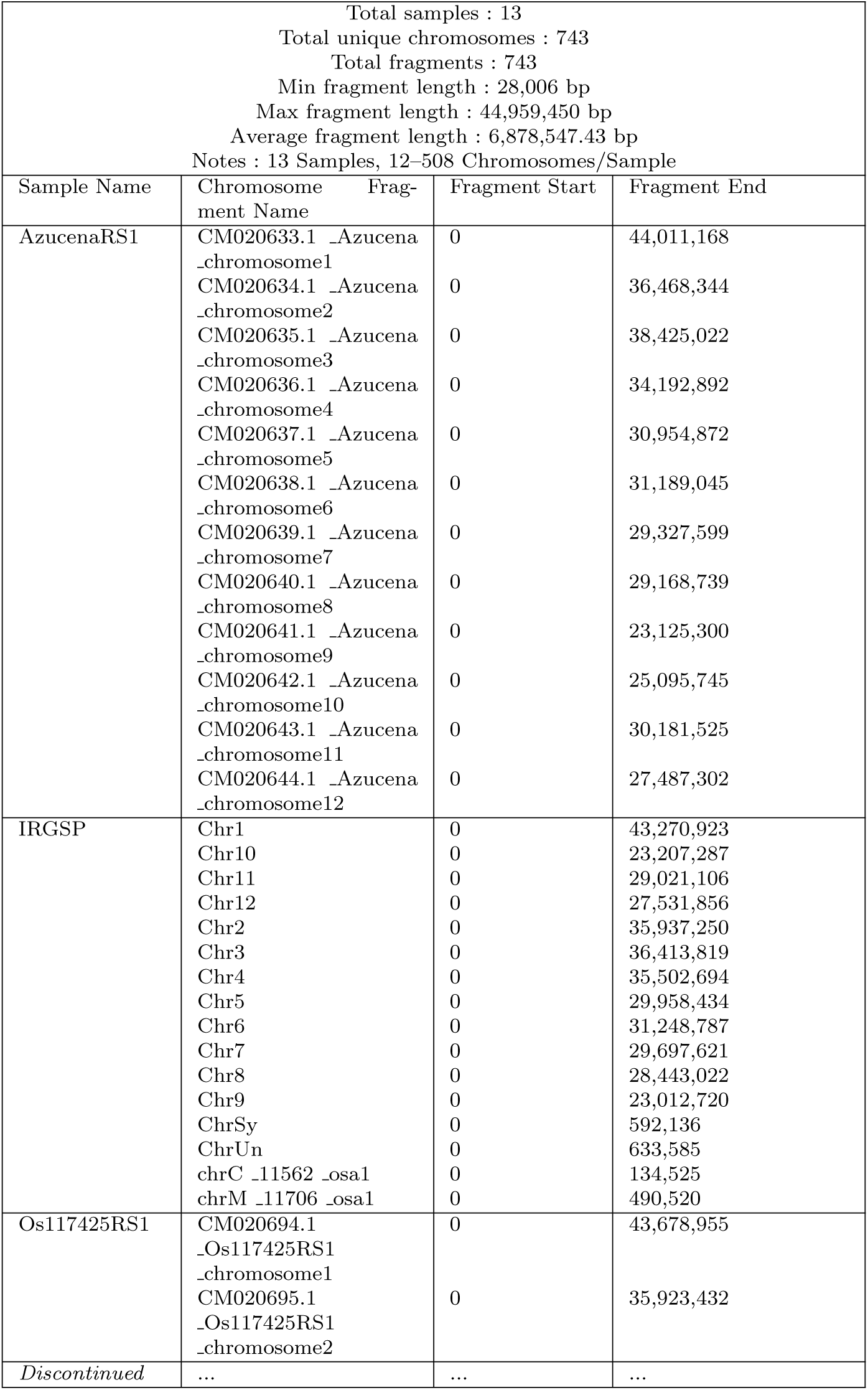
Adapted output that list (partially) of all chromosomes/contigs (called *Fragments*) from the embedded samples in the GFA from Asian rice, obtained through the gratools list chr --gfa NewRiceGraph MGC.gfa.gz --full command. The --full option provides the information of start and end of each Fragment. General statistics are included above as part of the same table content.

### 3.3 Extracting parts of a graph

GraTools can use the coordinate system from any of the embedded sample (haplotype) in the PVG, without having to re-import it, as it is the case in vg for instance. Indeed, the initial formatting of PVG under the GraTools data structure (import command) allows it to be able to recover any part of the graph based on any of these haplotypes, even if it has not been indicated as ”reference” during the PVG building or importing. In the following section, we exemplified the subgraph and FASTA extraction using two coordinate systems, one based on the reference used for building the PVG (here the NipponBare genome called IRGSP in this graph), the other one from another haplotype (namely IR64).

#### 3.3.1 Extracting a subgraph

While visualizing the whole PVG is quite unfeasible, having a view on a specific locus can help a lot understanding variations between haplotypes, using tools such as Bandage [34], as shown in Figure 2. The get subgraph will, as its name indicates, extract a subgraph based on the coordinates of *any* of the embedded path, not only based on the PVG reference coordinates. Here, using the Asian rice PVG from [15, 33], we extracted the Sub1 locus based on IR64 coordinates using the get subgraph command, the PVG having been build using the Nipponbare genome as reference. This locus harbors three genes (Figure 2), the Sub1B (yellow part in Figure 2) and Sub1C (green) that are present on all samples, and the Sub1A (in pink) that exists only in 4 samples upon 13, including IR64, and is responsible of the tolerance to submergence in Asian rice [4]. We used thus the IR64 coordinates to extract the subgraph for this main locus of more than 150 kb.

**Fig. 2.**
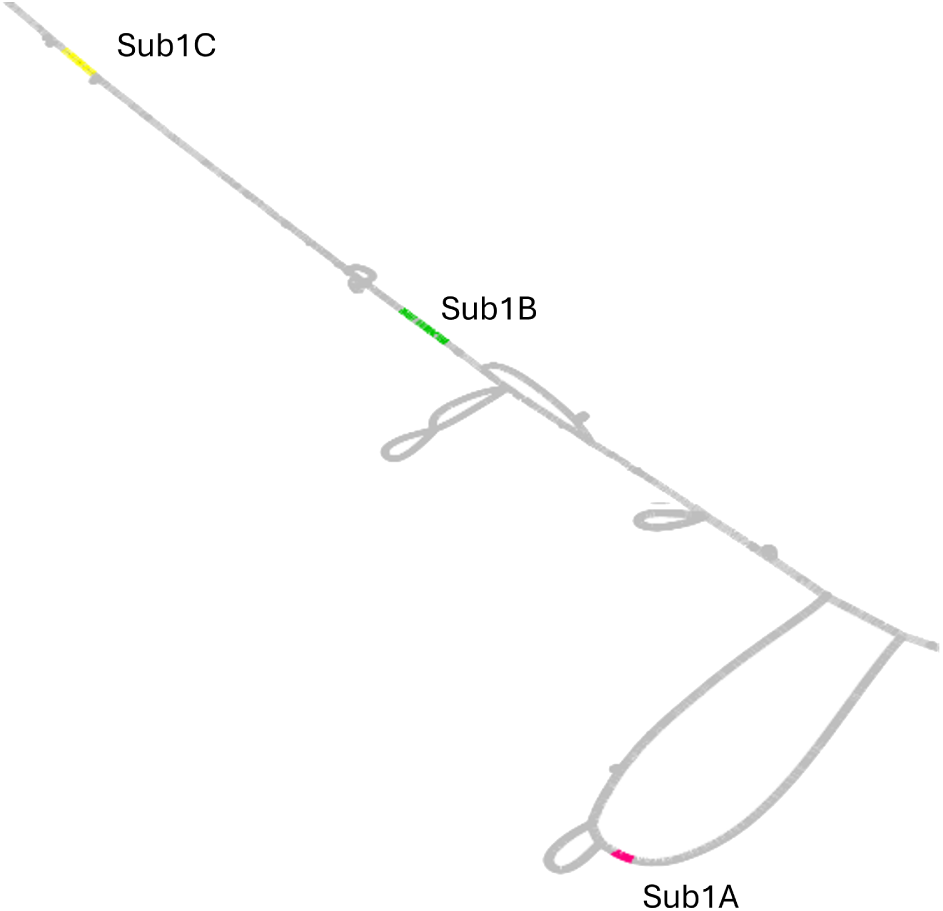
Bandage layout representation of the Sub1 locus in the Asian rice PVG from Marthe et al [15] extracted based on the IR64 coordinates. The 3 genes of the locus, namely Sub1C, Sub1B and Sub1A are depicted in yellow, green and pink, respectively on the graph structure. We used the following command: gratools get subgraph --gfa NewRiceGraph MGC.gfa.gz -sq OsIR64RS1 -chr CM020884.1 OsIR64RS1 chromosome9 --all-samples --start-query 7400227 --stop-query 7800003.

The extracted subgraph can contain the whole set of paths embedded in the graph (--all-samples option), but it can be restricted to a user-defined list of them. The node names in the newly extracted subgraph GFA file are the same as in the original GFA file, for a better forward analysis. In addition to a GFA file, GraTools can also output the corresponding FASTA sequences for each (required) path by adding the --build-fasta option. The same subgraph can also be extracted from e.g. the NipponBare reference genome using the specific coordinates and the following command: gratools get subgraph -g NewRiceGraph MGC.gfa.gz -sq IRGSP -chr Chr9 --start-query 6328847 --stop-query 6673971. These two graphs are identical (available as SuppData).

The time for extracting is partially dependent on the PVG depth (the number of embedded samples), as shown in Table 9. Indeed, extracting the same region in a larger graph is longer in CPU times (from less than one minute to 5) with a memory footprint increase (from 0.16 to 1.3GB) when going from a 20 samples PVG to a 69 one. However, it is still completely acceptable in terms of time.

**Table 9.**
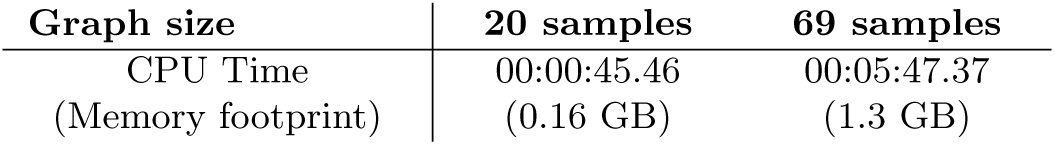
GraTools get subgraph command benchmarking. We tested to extract the same region (same coordinates on the Col0 reference genome), using a graph containing 20 samples and another with 69 samples for *A. thaliana*. The exact command for the 69 samples graph was gratools get subgraph -g 69 arabidopsis MGC.gfa -sq Col 0 -chr chr3 --all-samples -t 10 --start-query 8600000 --stop-query 8620000. The PVG were built as explained in Marthe et al [15] based on the sequences from Lian et al [40] and Kang et al [41].

#### 3.3.2 Recovering FASTA sequences from a subgraph/subregion

If only linear sequences for a given region are needed, the get fasta command extracts the corresponding FASTA sequences for all embedded genomes based on the coordinates of one of the embedded genome. As with the previous command, the output can be restricted to a specified list of paths. The provided coordinates may also be defined from a path that will not be included in the final output.

### 3.4 Analyses on the graph

#### 3.4.1 Core/Dispensable genome ratio

Most pangenomics analyses start with the estimation of the Core *vs*. Dispensable (or Pangenome) ratio. The pan ratio command computes this ratio, while allowing users to define the percentage of haplotypes harboring a node classified as Core (Table 10). For example, a strict core can be specified with 100% sharing (no missing haplotypes), while a more relaxed definition may require only 90% of haplotypes to share the node. In addition, the command can also report the proportion of specific sequences, defined as nodes present in only a small number of haplotypes. It also allows filtering of nodes according to their size (for instance, by computing the ratio only on structural variation larger than 50 bp, as shown in Table 10). In the current example here, for Asian rice, 61.81% of nodes larger than or equal to 50 bp (--fl option) are classified as core (with a fixed core at 90%), whereas only about 30% of nodes are considered core when no size filtering is applied. Indeed, the threshold of the minimal size of 50 bp removed 91.56% of nodes, indicating that the majority of nodes in the graph correspond to very small variant, typically SNPs.

**Table 10.** Core/Pangenome ratio statistics in the GFA from Asian rice, obtained through the gratools pan ratio -g NewRiceGraph MGC.gfa.gz --input-as-percentage -sm 90 -spm 10 -fl 50 command. Here the Core is computed as present in 90% or more of haplotypes, the specific present in 10% or less of haplotypes, with an interrogation of fragments with a length larger or equal to 50bp. The *Raw* value are not filtered (*i.e.* all nodes).

| Total segments in GFA: 26,461,214 |  |  |  |
| --- | --- | --- | --- |
| Total segments analyzed: 6,461,214 |  |  |  |
| Total segments passing length filter ($\geq 50$bp): 2,233,791 (8.44%) | | | |
| Core vs. Dispensable Segments — NewRiceGraph_MGC |  |  |  |
| Category | Count | Total | Percentage |
| Shared (Core) - Raw | 7,963,273 | 26,461,214 | 30.09% |
| Specific (Dispensable) - Raw | 3,824,526 | 26,461,214 | 14.45% |
| Shared (Core) - Filtered (Length $\geq 50$bp) | 1,380,679 | 2,233,791 | 61.81% |
| Specific (Dispensable) - Filtered (Length $\geq 50$bp) | 40,656 | 2,233,791 | 1.82% |
| Segments Filtered Out by Length | 24,227,423 | 26,461,214 | 91.56% |

Other commands linked to the depth of each node (meaning the number of haplotypes having this node in their path) exist in GraTools (Table 1), such as obtaining the statistics of node depths, or filtering out nodes with a minimal depth and size. All these commands can be restrained to a subset of embedded haplotypes, or excluding nodes under a specific threshold size.

#### 3.4.2 Recovering segments based on group specificity

The specific groups sample command proposes to extract nodes that are specific from a set of paths compared to another, or shared by a set, or absent in it. Here, as before, the term ”sample” refers then to a specific path from an haplotype in the PVG. It can be used for instance to recover nodes specific to a population compared to another (wild and cultivated, e.g.), or to find the nodes that are common to the whole population of a subspecies (or absent in it).

The Figure 3 shows an application in deciphering the number of nodes that are core-specific from each of the two rice subspecies (*indica* and *japonica*) in the Asian rice graph used in Marthe et al [15]. Here, we launched a suite of 3 simple commands to obtain the strict core (100% of the paths having the node) from different compartment in the PVG, as follows:

- gratools pan ratio -g NewRiceGraph MGC.gfa.gz --input-as-percentage -sm 100 -spm 10 provides the number of nodes shared by *all* samples (*Core shared*, Figure 3)
- gratools specific groups sample -g NewRiceGraph MGC.gfa.gz -sla samples indica -slb samples japonica provides the *indica* core, *i.e.* the number of nodes present in *all indica* but *no japonica* (*Core Indica*, Figure 3)
- gratools specific groups sample -g NewRiceGraph MGC.gfa.gz -sla samples japonica -slb samples indica provides *japonica* core, meaning the number of nodes present in *all japonica* but *no indica* (*Core Japonica*, Figure 3)

**Fig. 3.**
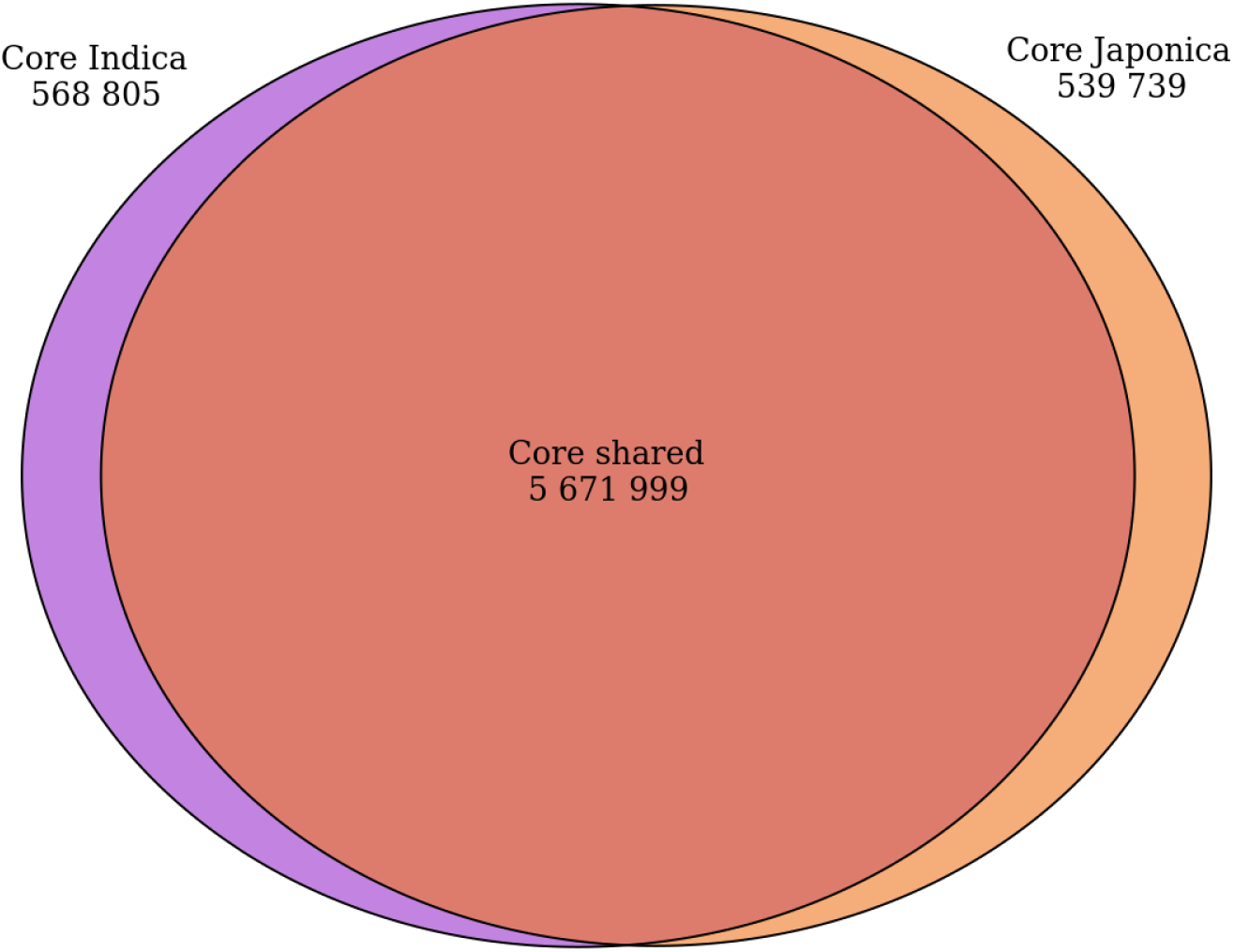
Venn diagram representing the *indica* and *japonica* subspecies specific strict core nodes (shared by all samples) in the Asian Rice PVG from Marthe et al [15], computed using specific groups sample and ratio commands. For each subspecies (purple for the *indica*, and orange for the *japonica*), we indicate the total number of nodes shared by all samples. The intersection (Core shared) represent nodes shared by 100% of the haplotypes of the Asian rice PVG.

#### 3.4.3 Advanced usage of the GraTools data structure

While GraTools proposes a large panel of command-line analyses, it cannot cover all possible use cases. However, as its internal files are BAM and BED format, they can be directly reused for more advanced and specific analyses. Here, we present an example of such an analysis, which requires higher bioinformatics proficiencies than the previous analyses. We were interested in knowing the exact size of the pangenome, both in node number and total sequence length, for each of the two rice subspecies. All the codes used in this analysis are referenced in the ”Availability of data” section 5.

We started by launching a series of specific groups sample command, one per sample, to identify nodes present in this individual but absent from all samples belonging to the opposite group. We then identified nodes present in at least one of the 8 *indica* but absent from all 5 *japonica*, and vice-versa. We then mixed the *indica* nodes to obtain an *indica*-only node list, and do the same for the *japonica*. Once these specific *indica* and *japonica* node lists obtained, we summed their size using a custom python script and the pysam API upon the BAM file containing the nodes information (see section 5 for the detailed commands). We then summarized the results as a Venn diagram, figured out in Figure 4.

**Fig. 4.**
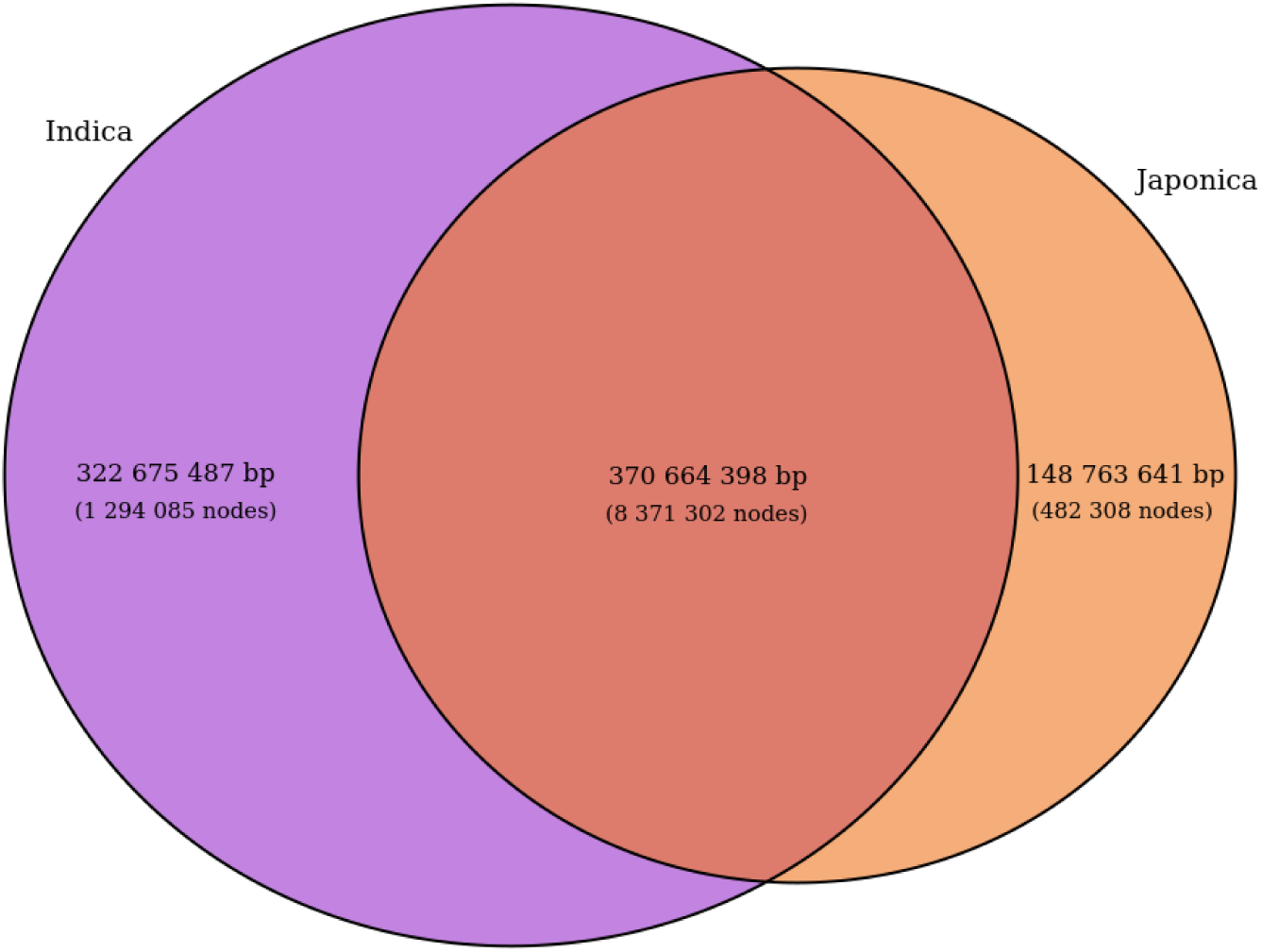
Venn diagram representing the *indica* and *japonica* subspecies specific nodes in the Asian Rice graph from Marthe et al [15], computed using the specific groups sample command (see text for more details). For each subspecies (purple for the *indica*, and orange for the *japonica*), we indicate the sum in bp of the total nodes (specific or shared).

As expected, the *indica* (8 samples upon 13) specific part is larger than the *japonica* (5 samples) one, as discussed in Zhou et al [33]. In Asian rice, the expected genome size is of 380 to 410 Mb, and here the core part represent roughly 90-95% of the genome of each individual (371 Mb). The mean node size in the core genome is 44 bp, whereas it reaches 249 and 308 bp for *indica* and *japonica* part respectively. This indicate that most variations distinguishing the two groups correspond to larger structural variations rather than SNP, as seen in Qin et al [35].

## 4 Discussion

Obtaining a PVG has become a classical step in the study of genomic variations across species ([9, 36]). However, once the PVG is constructed, structural variants (SV) are often extracted from the graph (mostly using the vg deconstruct command) to obtain a classical VCF file, capturing only a subset of the complex, nested SVs ([37]). Consequently, analyses frequently came back to linear VCF-based representation due to the lack of user-friendly tools that can directly exploit PVGs to answer different biological questions. To overcome these limitations, we developed GraTools to meet most of the needs of end-users with basic to advanced knowledge in pangenomics and in command line usage. GraTools enables a rapid and innovative analysis directly on PVGs, including the computation of core and dispensable genome ratios, estimation of node depth across individuals, and extraction of sequences from genomic coordinates defined on any genome embedded in the graph. All these analyses are performed rapidly (e.g. get subgraph command, see Table 9) once the import in GraTools format is done.

As GFA files are complex, huge tabular text-files difficult to manipulate natively, GraTools, similarly to vg [8] and ODGI [13], import the data contained in the GFA in dedicated optimized files. However, GraTools commands always refer to the GFA file itself as input, and not to these optimized files: it ensures a transparency and efficiency of usage for biologists. Moreover, the import command is intrinsic to any of the other commands: if it has not yet been executed and the BAM/BED files obtained when asking for another command (e.g. gratools stats), it will import the data in BAM and BED format automatically. In terms of performance for importing, GraTools is still indeed slower compared to vg and ODGI (see Table 4 and 3.1 part). Being aware of the limits we observed nevertheless in the GraTools performance for importing, we plan to refactor this part in the near future, allowing to save time and space, and homogenizing all our ”indexing” approaches between our tools from the GraSuite (such as Savanache [38]). However, GraTools is the only tool that does not require a new indexing or any complex file manipulation when changing the system of coordinates, and thus is more efficient long term. Indeed, as shown in 3.3 and 3.3.2 sections, users can immediately change the path they use for providing coordinates, without having to re-import the whole data based on this ”new reference”. In addition, all these extractions refer to the same node name system, from the initial GFA file to all the extraction, ensuring a very efficient backward compatibility.

The GraTools commands (Table 1) are quite intuitive and stream-lined, compared to the multiple commands to perform for the other tools to obtain the same results (Table 2). They also permit to obtain rapidly information on the PVG and on the haplotypes embedded in (see 3.2.1 and 3.2 sections), extracting subgraphs (3.3.1 section), or to have a biological insight on their data (as for the Core/Dispensable analysis on Asian rice, 3.4.1 section). Finally, for power-users, the BAM and BED transformation of the GFA file give access to large results quite fast (3.4.3). In this last case, we were able to obtain the data in less than one hour without any optimization of codes, much faster than the original analysis from Zhou et al [33], but with the same results.

## 5 Conclusion

In the present paper, we present GraTools, an intuitive and user-friendly command line tool to manipulate, extract and analyze pangenome variation graphs in GFA format and member of the GraSuite [1]. It has been tested on PVGs from various origins and species, and has been shown to be fast and efficient. It was designed with in mind the use of GFA for subsequent analyses, in particular population genetics or evolution ones, but can also be used for breeding (detection of specific nodes in a set of elites lines e.g.), genomic medicine (detection of missing/additional nodes in population with genetic illness) or even antibiotic resistance analyses (detection of additional segments in resistant strains). As a large set of commands are included within GraTools, users can manipulate their GFA within a single environment, and re-use it fast due to the conservation of the intermediate files and indexes in their user space (or shared one). The outputs, messages and log files are human-readable and understandable, allowing users to manage them easily. Thus, GraTools streamlines the manipulation of pangenome variation graph in GFA format, allowing to really unleash the potential of pangenome graphs in higher eukaryotes.

## Supplementary information

All Supplementary data are available at https://dataverse.ird.fr/dataverse/gratools.

## Acknowledgments

The authors acknowledge the ISO 9001 certified IRD i-Trop HPC at IRD Montpellier for providing HPC resources that have contributed to the research results reported within this paper. URL: https://bioinfo.ird.fr/. The analyses were also performed on the Core Cluster of the Institut Fraņcais de Bioinformatique (IFB) (ANR-11-INBS-0013).

## Declarations

- Funding: Nina Marthe is supported by a PhD funding from the Agence Nationale de la Recherche as part of the France2030 program under the reference ”ANR-22PEAE-4”. Camille Carrette is supported by a CIFRE ANRT between Syngenta Seeds SAS and IRD. Mourdas Mohamed is supported by a PlantAlliance/SofiProteol/IRD joint project.
- Conflict of interest/Competing interests: None
- Ethics approval and consent to participate: Not applicable
- Consent for publication: Yes
- Author contribution: CTD, FS and SR managed the project; SR, NM, MM, CC and CTD implemented the code; SR, FS and all authors developed the algorithms; CTD, FS and SR drafted the manuscript; all authors corrected and validated the manuscript.

## Availability of data and materials

The rice pangenome graph used for figures and analyses is available in Marthe et al [15]

- Data availability: https://dataverse.ird.fr/dataverse/gratools
- Code availability: https://forge.ird.fr/diade/gratools for the most recent version, and https://dataverse.ird.fr/dataverse/gratools for the version used in the current manuscript

## Availability and Requirements

- Project name: GraTools
- Project homepage: https://gratools.readthedocs.io/
- Operating system(s): Linux, MacOS, Windows (with WSL)
- Programming language: Python 3.12+
- Other requirements: Bedtools
- License: GNU GPLv3

